# An immune signature of postoperative cognitive decline in elderly patients

**DOI:** 10.1101/2024.03.02.582845

**Authors:** Franck Verdonk, Amélie Cambriel, Julien Hedou, Ed Ganio, Grégoire Bellan, Dyani Gaudilliere, Jakob Einhaus, Maximilian Sabayev, Ina A. Stelzer, Dorien Feyaerts, Adam T. Bonham, Kazuo Ando, Benjamin Choisy, David Drover, Boris Heifets, Fabrice Chretien, Nima Aghaeepour, Martin S. Angst, Serge Molliex, Tarek Sharshar, Raphael Gaillard, Brice Gaudilliere

## Abstract

Postoperative cognitive decline (POCD) is the predominant complication affecting elderly patients following major surgery, yet its prediction and prevention remain challenging. Understanding biological processes underlying the pathogenesis of POCD is essential for identifying mechanistic biomarkers to advance diagnostics and therapeutics. This longitudinal study involving 26 elderly patients undergoing orthopedic surgery aimed to characterize the impact of peripheral immune cell responses to surgical trauma on POCD. Trajectory analyses of single-cell mass cytometry data highlighted early JAK/STAT signaling exacerbation and diminished MyD88 signaling post-surgery in patients who developed POCD. Further analyses integrating single-cell and plasma proteomic data collected before surgery with clinical variables yielded a sparse predictive model that accurately identified patients who would develop POCD (AUC = 0.80). The resulting POCD immune signature included one plasma protein and ten immune cell features, offering a concise list of biomarker candidates for developing point-of-care prognostic tests to personalize perioperative management of at-risk patients. The code and the data are documented and available at https://github.com/gregbellan/POCD.

**Teaser:** Modeling immune cell responses and plasma proteomic data predicts postoperative cognitive decline.

## Introduction

Annually, patients aged 60 and older undergo over 90 million major surgical procedures worldwide(*1*), a figure projected to quadruple within the next four decades(*2*). Postoperative cognitive decline (POCD) represents a significant challenge in this population, leading to impairments in verbal and visual memory, language comprehension, visuospatial abstraction, attention, and concentration following surgery. It ranks among the most prevalent complications after major surgery, affecting 25% to 55% of elderly patients for up to a year(*3*, *4*). POCD correlates with increased mortality, earlier retirement, and heightened reliance on social financial assistance(*5*, *6*). Although there is currently no specific treatment for POCD, a preventive approach including cognitive prehabilitation training, physical exercise, and preoperative geriatric consultation has demonstrated efficacy in mitigating POCD risk(*7–9*). However, estimating an individual patient’s POCD risk remains challenging as previous attempts relying on clinical factors have yielded limited predictive power(*10*), severely impeding implementation of preventative interventions. Accurate biological or clinical biomarkers of POCD are urgently needed to develop a prognostic test to guide personalized management of elderly surgical patients(*11*).

Emerging preclinical evidence suggests the systemic immune response to surgical trauma induces sustained microglial activation(*12–14*), resulting in key pathobiological processes driving POCD, including synaptic dysfunction, inhibition of neurogenesis, and neuronal apoptosis(*15*). In response to surgery, trauma-induced circulatory mediators, including high mobility group box (HMGB)-1, heat shock proteins, and toll-like receptors (TLR), activate the inflammasome pathway in innate immune cells, resulting in systemic pro-inflammatory cytokine release(*16–18*). Several of these cytokines, such as interleukin (IL)-1β(*19*), tumor necrosis factor (TNF)-α(*20*), or IL-6(*21*) have been identified as causative factors of POCD in animal models. In rats, depleting bone marrow-derived macrophages disrupts the innate immune response to surgery and reduces postoperative memory dysfunction, highlighting a critical role of the myeloid phagocytic system in the pathophysiology of POCD(*22*). Despite important insights from preclinical studies into the neuro-inflammatory mechanisms of POCD, human research in patients undergoing surgery is lacking. Prior clinical studies focusing on biomarkers of neuronal damage(*23*) or systemic inflammation, including C-reactive protein and IL-6(*24*), have only revealed weak biological correlates of POCD. Given the dynamic and multicellular nature of the surgical immune response, prior analyses may have overlooked strong signals, as specific immune cell phenotype or functional responses were not fully examined.

Highly multiplexed single-cell proteomic technologies, such as mass cytometry, provide powerful platforms to characterize complex and multicellular inflammatory processes like the human immune response to surgery(*25–29*). Further integration of single-cell and plasma-based proteomic modalities allows interrogation of multiple interconnected biological systems and otherwise unrecognized pathophysiological crosstalk. Merging multiple omic modalities can reveal biomarkers from several biological domains that, together, provide higher power to predict perioperative outcomes(*30–33*).

Here, we employed an integrated approach combining the functional analysis of circulating immune cell subsets via mass cytometry and multiplex assessment of plasma proteins to quantify dynamic immunological changes in elderly patients before and after major orthopedic surgery. The primary aim was to provide a comprehensive analysis of innate and adaptive immune cell trajectories differentiating patients with and without POCD. A secondary goal was to determine whether integrating single-cell and plasma proteomics with clinical data collected before surgery would yield a robust predictive model identifying patients at risk for POCD.

## Results

### Patient characteristics

Thirty-three patients (aged 72.8±7.1 years) scheduled for major orthopedic surgery were enrolled in the study. Of these, 26, including 20 females (76.9%), underwent cognitive assessment seven days after surgery and were included in the analysis (see CONSORT flow chart **Fig. S1**), 11 (42.3%) of whom had developed POCD (**Fig. 1**). Amongst patients with POCD at postoperative day (POD)7, four (36.3%) presented with persistent POCD at POD90, six (54.5%) had recovered and one did not undergo cognitive evaluation. Patient characteristics are described in **Table 1**. Patients with and without POCD did not differ in terms of known clinical risk factors for POCD such as age, history of degenerative or psychiatric disease, or education level(*34*). The type of surgery or anesthesia including peri-operative administration of ketamine or dexamethasone did not differ between the two patient groups (**Table S1**). Postoperative follow-up assessments, including postoperative depression (Hospital Anxiety and Depression Scale score ≥ 8), pain levels, and morphine consumption, revealed no differences between the two patient groups (**Table S2**).

**Figure 1:**
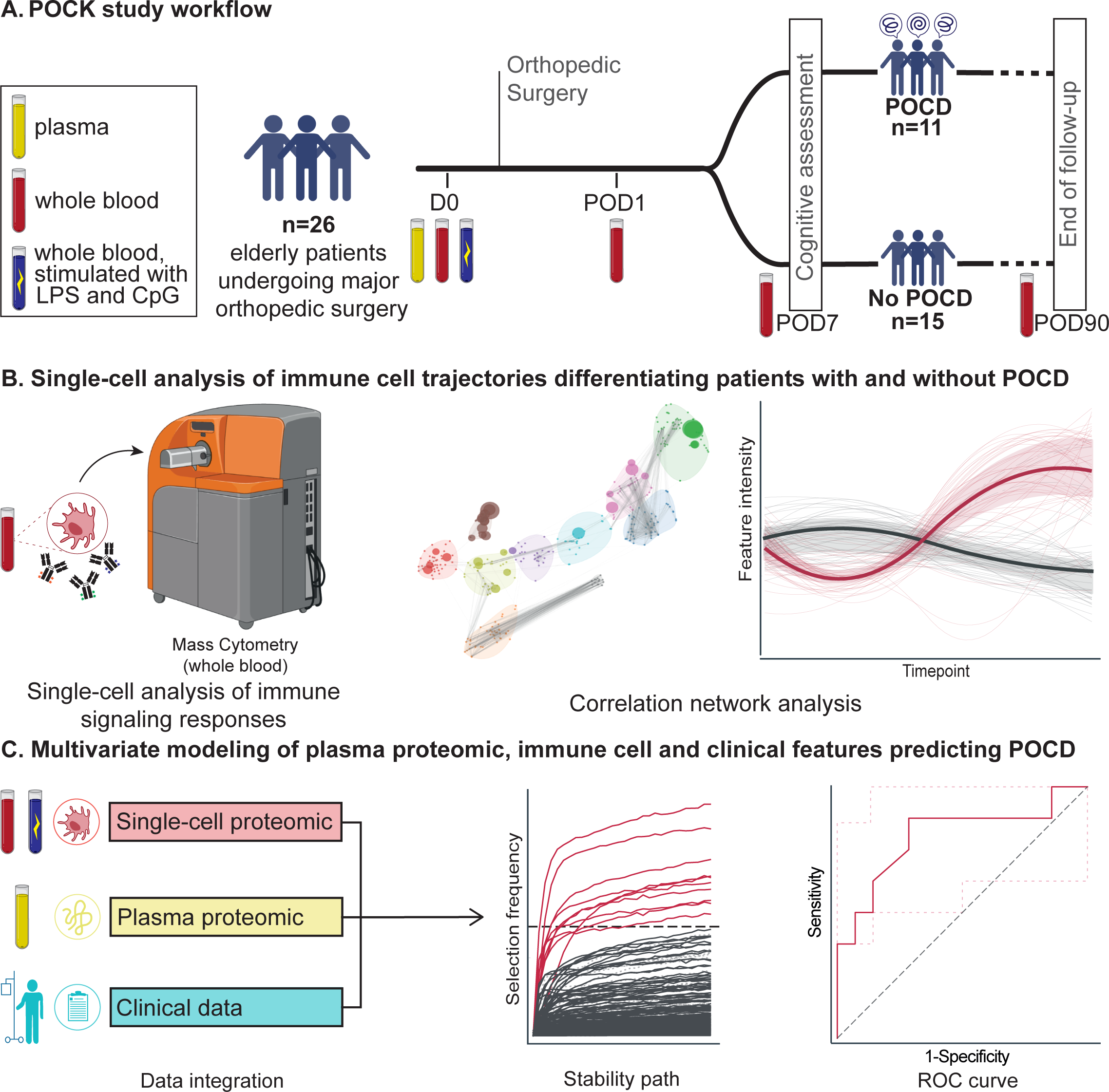
Study design for combined single-cell and plasma proteomic immune profiling of post-operative cognitive decline (POCD). **A**. Peripheral blood samples were collected from 26 patients undergoing primary hip replacement surgery at four timepoints: before surgery at day (D)0 and on postoperative day (POD)1, POD7, and POD90. Among them, 11 patients developed POCD seven days after surgery (POCD group), while 15 did not (control group). **B.** Longitudinal analysis of immune cell trajectories differentiating the POCD and control groups was performed using single-cell mass cytometry. **C.** In addition, a multivariable predictive modeling approach integrating the plasma proteomic (SOMAscan manual assay), single-cell proteomic (cytometry by time of flight mass spectrometry, CyTOF), and clinical datasets obtained before surgery (D0) was employed to identify preoperative biomarker candidates predictive of POCD. *Abbreviations: LPS: lipopolysaccharide*

**Table 1:** Clinical and demographic characteristics of the study population at baseline. No significant differences were observed between the two groups with respect to any known risk factors for POCD measured in the study. Values are expressed as numbers (percentage %), means (standard deviation) or medians [IQR]. Mann-Whitney U tests were performed. *Abbreviations: HADS: Hospital Anxiety and Depression Scale; Hb: Hemoglobin; IQR: interquartile range, MoCA; Montreal Cognitive Assessment score; POCD: Postoperative Cognitive Decline, SD: Standard deviation*

| Characteristics |  | Whole<br>population<br>(n=26) | No POCD<br>(n=15) | POCD<br>(n=11) | p-value |
| --- | --- | --- | --- | --- | --- |
| Sex, female |  | 20 (76.92%) | 13 (86.67%) | 7 (63.64%) | 0.348 |
| Age, mean (SD) |  | 72.88 (7.1) | 72.2 (6.27) | 73.82 (8.33) | 0.595 |
| Education | University or higher | 13 (52%) | 8 (57.14%) | 5 (45.45%) | 0.871 |
|  | High school | 8 (32%) | 4 (28.57%) | 4 (36.36%) |  |
| Charlson comorbidity index, Median [IQR] |  | 3.5 [3;5] | 4 [3;5] | 3 [2.5;4.5] | 0.825 |
| Obstructive sleep apnea |  | 3 (12.64%) | 1 (7.66%) | 2 (18.18%) | 0.556 |
| Hypertension |  | 14 (53.85%) | 7 (46.67%) | 7 (63.64%) | 0.453 |
| History of neurodegenerative or psychiatric disease |  | 4 (15.38%) | 2 (13.33%) | 2 (18.18%) | 1 |
| Current smoker |  | 1 (3.85%) | 0 (0%) | 1 (9.09%) | 0.423 |
| Preoperative anemia<br>(Hb <13 g.dl<sup>-1</sup> [male], Hb <12 g.dl<sup>-1</sup> [female]) |  | 1 (3.85%) | 0 (0%) | 1 (9.09%) | 0.423 |
| Cognitive, anxiety and depression status |  |  |  |  |  |
| Montreal Cognitive Assessment score, Median [IQR] |  | 25 [22;27] | 23 [20;27] | 26 [22.5;27] | 0.291 |
| MOCA range | [20-26] (mild cognitive impairment) | 14 (53.85%) | 7 (46.67%) | 7 (63.64%) | 0.195 |
|  | <20 (severe cognitive impairment) | 4 (15.38%) | 4 (26.67%) | 0 (0%) |  |
| Anxiety (HADS Anxiety score ≥8) |  | 13 (50%) | 8 (53.33%) | 5 (45.45%) | 0.691 |
| Depression (HADS Depression score ≥8) |  | 4 (16%) | 2 (14.29%) | 2 (18.18%) | 1 |
| Type of surgery | Shoulder arthroplasty | 1 (3.85%) | 0 (0%) | 1 (9.09%) | 0.964 |
|  | Spine surgery | 5 (19.23%) | 2 (13.33%) | 3 (27.27%) |  |
|  | Total hip arthroplasty | 11 (42.31%) | 6 (40%) | 5 (45.45%) |  |
|  | Total knee arthroplasty | 3 (11.54%) | 2 (13.33%) | 1 (9.09%) |  |
|  | Revision total hip arthroplasty | 6 (23.08%) | 4 (26.67%) | 2 (18.18%) |  |
| Type of anesthesia | General | 18 (69.23%) | 9 (60%) | 9 (81.82%) | 0.395 |
|  | General and regional (i.e., peripheral nerve block) | 8 (30.77%) | 6 (40%) | 2 (18.18%) |  |

### Longitudinal assessment of perioperative immune cell distribution and signaling states

Whole blood samples collected longitudinally before (day (D)0, day of surgery) and after surgery (postoperative day (POD)1, POD7, and POD90) were analyzed using a 39-parameter mass cytometry immunoassay for single-cell profiling of peripheral immune cell distribution and intracellular signaling protein phosphorylation states. The single-cell mass cytometry dataset was visualized using a uniform manifold approximation and projection (UMAP) layout highlighting the dynamic changes in signaling states of distinct immune cell compartments at each of the perioperative timepoints (**Fig. 2A**). In accordance with previous mass cytometry studies in patients undergoing surgery(*35*), the UMAP analysis highlighted canonical trauma-induced immune cell responses, including the rapid phosphorylation of signal transducer and activator of transcription (STAT)3 (pSTAT3) at POD1 in both innate (monocyte, neutrophil) and adaptive (CD4^+^ and CD8^+^ T cell) subsets (**Fig. 2B**) and the biphasic response (de-phosphorylation followed by phosphorylation) of key elements of the myeloid differentiation primary response 88 (MyD88) signaling pathway in innate immune cells, including (p)P38, (p)ERK1/2, (p)MAPKAPK2, and (p)NF-κB (**Fig. 2B, Fig S2**).

**Figure 2:**
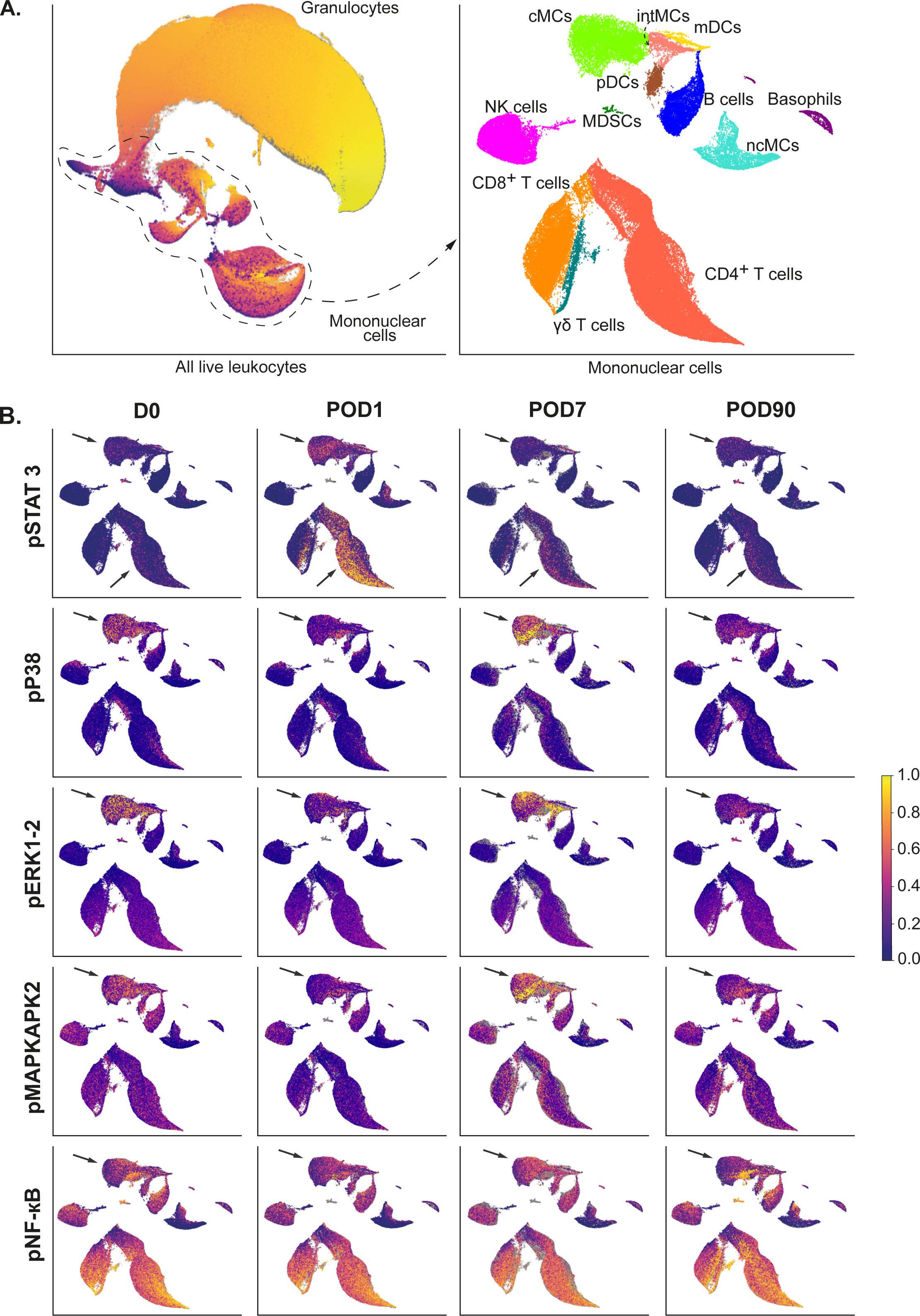
Longitudinal analysis of immune cell events before and after surgery with mass cytometry reveals cell-type and signaling-specific responses. **A**. Uniform manifold approximation and projection (UMAP) representation of the single-cell mass cytometry dataset. *Upper panel*: all live leukocytes, including neutrophils and mononuclear cells; *lower panel*: UMAP representation of mononuclear cells only. UMAPS are clustered by cell types and annotated. **B.** UMAPs representing all mononuclear cells colored according to intra-cellular signaling at D0 (before surgery), POD1, POD7, and POD90. *Abbreviations: cMCs: classical monocytes; CREB: cAMP-response element binding protein;* ^γδ^*T cells: gamma delta T cells; I*κ*B: inhibitor of* κ*B; intMCs: intermediate monocytes; mDCs: myeloid dendritic cells; MDSCs: myeloid-derived suppressor cells; NF-*κ*B: Nuclear factor-*κ*B; ncMCs: non-classical monocytes; NK: natural killer; pDCs: plasmacytoid dendritic cells; p: phosphorylated; JAK/STAT: Janus kinase/signal transducer and activator of transcription; rpS6: ribosomal protein S6*.

To complement the single-cell UMAP visualization, immune cell subsets were identified using an established gating strategy(*26*) (**Fig. S2**), and a total of 488 immune features were quantified, corresponding to either the frequency or the intracellular signaling activity of 11 proteins (measured as protein phosphorylation) within 31 innate and adaptive immune cell subsets. To examine the frequency and signaling trajectories of gated immune cell subsets in response to surgery, the high-dimensional immunological dataset was visualized on a correlation network that emphasized the interconnectivity of innate and adaptive immune features after surgery over time (**Fig. 3**). An unsupervised analysis using k-means clustering identified ten meta-clusters of highly correlated immune features (**Fig. 3A**, **Fig. S4,** see methods). Immune features within each meta-cluster evolved synchronously over time, likely reflecting the integration of multiple extracellular signals released in the circulation in response to surgical trauma (including alarmins and inflammatory cytokines) into synchronized immune cell trajectories (**Fig. 3B**). Meta-clusters 3, 5, 7, and 8 predominantly contained signaling response features that pointed at the JAK/STAT signaling pathways (98%, 100%, 100%, and 68% respectively). Other meta-clusters (e.g., 0, 2, 6, and 9) were mainly composed of signaling response features within the MyD88 signaling pathways (100%, 54%, 87%, and 80% respectively, **Table S3**).

**Figure 3:**
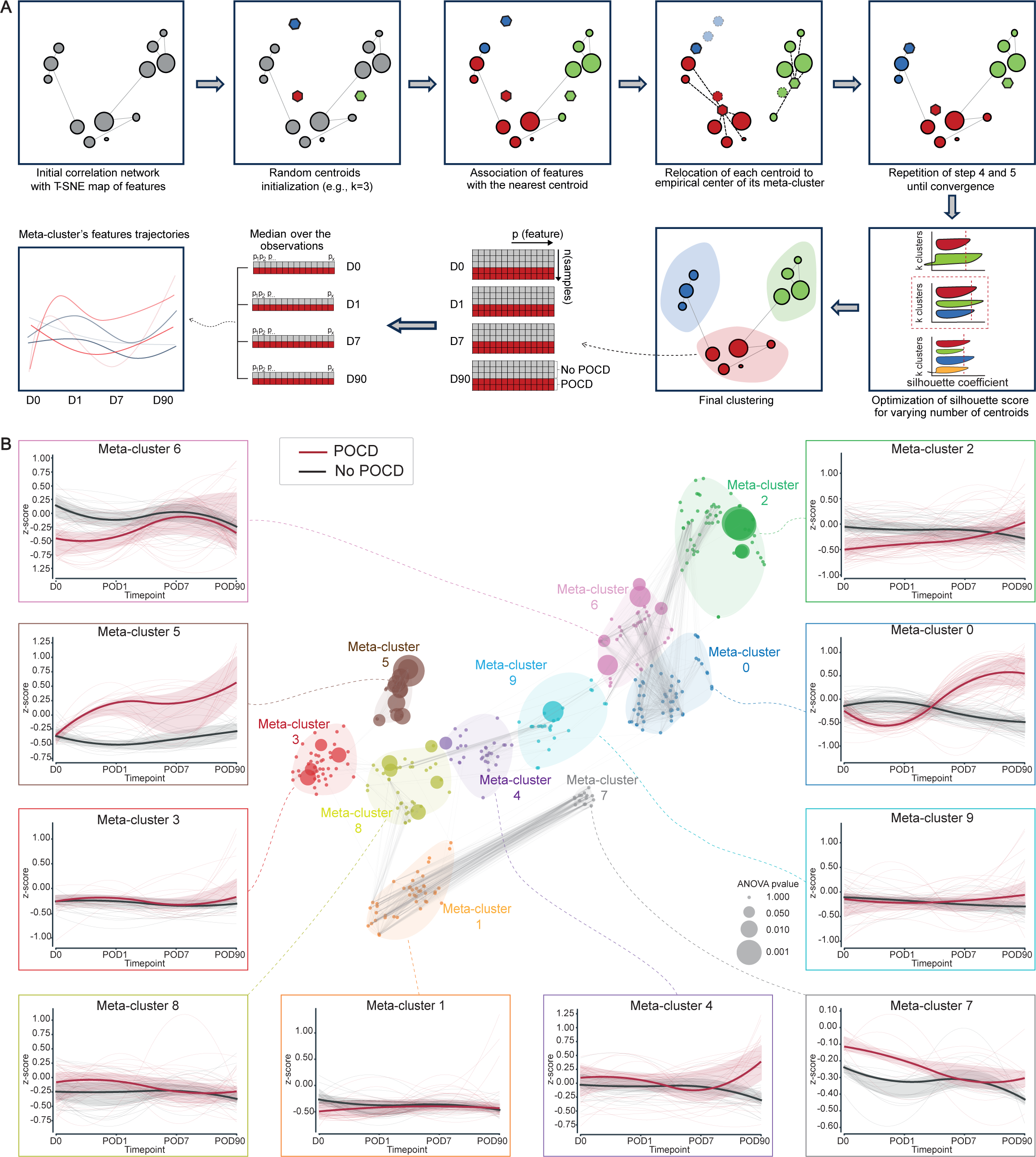
Longitudinal analysis identifies immune cell trajectories differentiating patients who later developed POCD from controls. **A**. Identification of immune cell feature clusters behaving synchronously in response to surgery (meta-clusters): An original dataset is constructed with a size of 4*n* x *p*, where “*n*” represents the 26 patients included in the study, and “*p*” represents the number of mass-cytometry features measured at each of the four timepoints (D0, POD1, POD7, and POD90). A two-dimensional embedding map is generated via t-SNE. An empirical determination of a number (k) of meta-clusters, each containing immune cell features with similar time-dependent trajectories, is achieved using an iterative nearest centroid approach (Fig. S3 and methods). After establishing the final meta-clusters, meta-cluster trajectories are extracted by calculating the median value of features within each meta-cluster. **B.** *Center panel.* A correlation network illustrates mass cytometry immune cell features and meta-clusters. Each node represents an immune cell feature, e.g., the frequency or median signaling activity of a specific immune cell subset. The size of each node corresponds to the p-value of the main effect of POCD calculated from an ANOVA. Edges represent a correlation coefficient (R)>0.7 between two features. Meta-clusters are encircled. *Peripheral panels*: For each meta-cluster, meta-cluster trajectories are displayed as red/black splines for the POCD and the control groups, respectively, with shaded interquartile ranges. Features are z-scored by subtracting the empirical mean of the feature and dividing it by the empirical standard deviation for each value.

Overall, results from the trajectory analysis were consistent with prior mass cytometry studies(*25*, *36*) and highlighted the remarkably orchestrated organization of immune cell responses to surgical injury. Interestingly, while the temporal behavior of immune cell responses was generally conserved across patients, we observed significant inter-patient variability in the amplitude of these immune cell trajectories, prompting us to investigate whether this variability reflected patient-specific differences associated with POCD.

### Single-cell immune response trajectories differentiate patients with and without POCD

To determine whether postoperative immune cell trajectories differed between patients with and without POCD, we compared the median of all features within a given meta-cluster for each group using a two-way analysis of variance (ANOVA) with fixed effect analysis of cognition and time relative to surgery. The analysis identified time-dependent differences between the POCD and control groups for meta-clusters 0, 5, 6, and 7 (p-values = 7×10^-26^, 6×10^-19^, 1×10^-8^ and 8×10^-11^, respectively). These meta-clusters also exhibited the strongest sensitivity to temporal progression (**Fig. 3B**), providing a meta-cluster-level synopsis of immune cell trajectories diverging between patients with and without POCD after surgery. The results are reminiscent of recent transcriptomic, proteomic, and flow cytometry analyses(*27–29*, *37*, *38*) linking immune trajectories with clinical recovery after surgery and emphasize the importance of studying immune responses dynamically as clinically-relevant biological differences could otherwise go undetected(*39*). Immune cell meta-clusters can be viewed as communities of synchronous immune features that can provide further information about immune cell-or signaling-specific states associated with POCD. Immune features within meta-clusters differing the most between the two patient groups were examined in further detail.

### JAK/STAT signaling trajectories diverge early after surgery in patients who develop POCD

One of the most prominent differences between the two patient groups was the endogenous pSTAT3 signal in adaptive immune cell compartments, including in CD4^+^ T cell subsets (naïve and memory) and regulatory T cells (meta-cluster 5, **Fig. 4A**). Specifically, the pSTAT3 signal within these adaptive cell subsets diverged as early as POD1 in patients with POCD and remained higher than in the non-POCD group throughout all postoperative timepoints (p = 0.005, 0.008, and 0.012 respectively, ANOVA with fixed effect for group). Fixed effect analysis of the time variable indicated that observed pSTAT3 signal differences in these adaptive immune cell subsets were also time-dependent (p = 5×10^-16^, 2×10^-11^, and 2×10^-10^ respectively, ANOVA, fixed effect for time). Additional differences in pSTAT3 were observed in innate immune cells, including neutrophils, classical monocytes (cMCs), non-classical (nc)MCs, and intermediate MCs (p = 0.014, 0.011, 0.038, and 0.047 respectively, ANOVA, fixed effect for group, **Table S4**). In both innate and adaptive immune compartments, the pSTAT3 signal did not return to baseline at POD90, suggesting prolonged innate and adaptative immune dysfunction associated with POCD after surgery. Together, these results suggest JAK/STAT signaling responses in both innate and adaptive cells are exacerbated as early as POD1 in patients who develop POCD, and that this activated state can extend at least 90 days after surgery(*35*).

**Figure 4:**
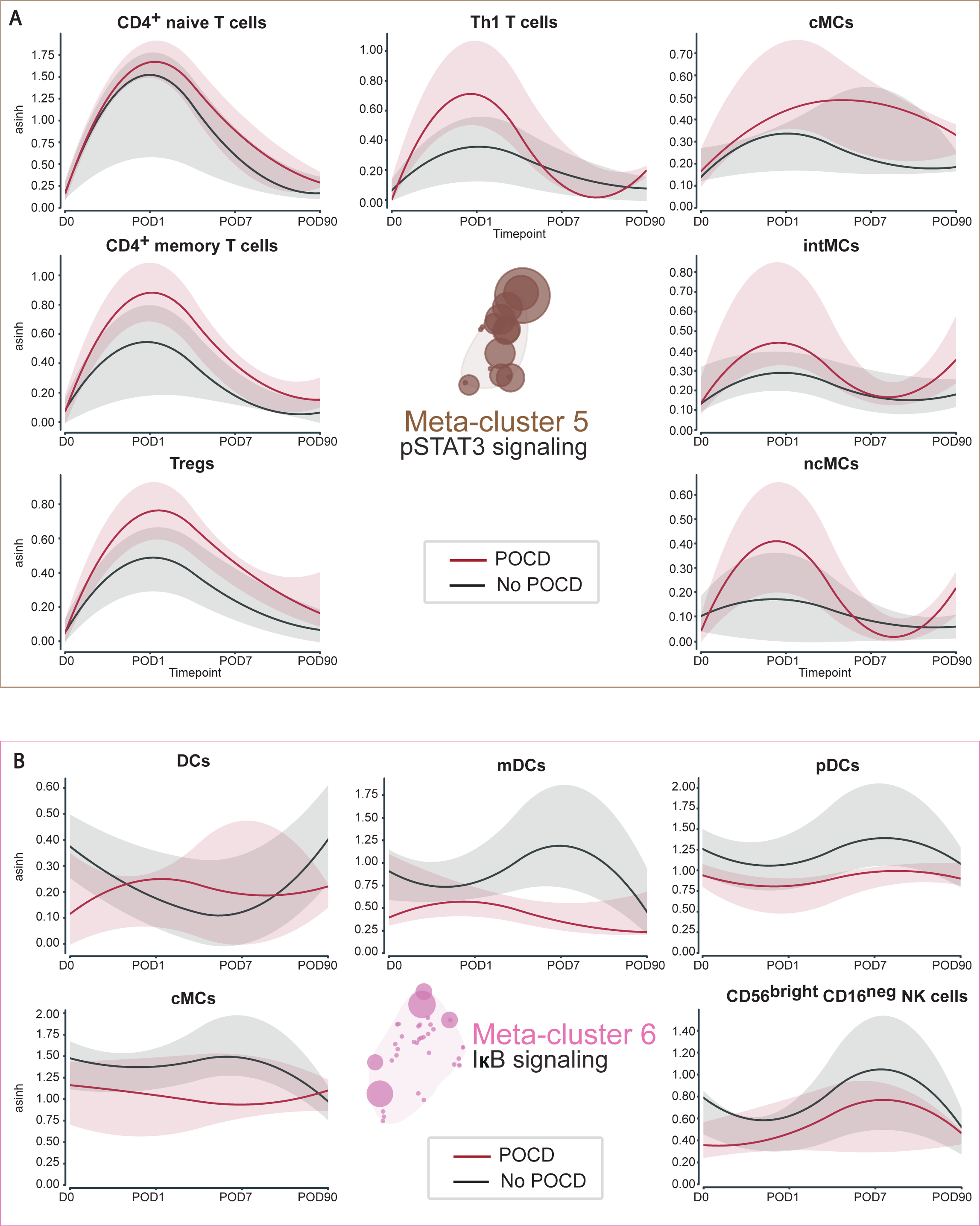
JAK/STAT and MyD88 immune trajectories differentiate patients with and without POCD. **A**. Immune cell features contained within meta-cluster 5. Median (splines) and interquartile range (shaded area) expression of the basal pSTAT3 signal (arcsinh transformed) in CD4^+^ naïve T cells, CD4^+^ memory T cells, regulatory T cells (Tregs), type 1 T helper (T_h_1) cells, cMCs, ncMCs, intMCs at D0, POD1, POD7, and POD90 in the POCD (red) and control (black) groups. **B.** Immune cell features contained within meta-cluster 6. Median (splines) and interquartile range (shaded area) expression of the basal pIκB in DCs, cMCs, pDCs, mDCs and CD56^bright^CD16^neg^ NK cells, in the POCD (red) and control (black) groups. *Abbreviations: ncMCs: non-classical monocytes; pDCs: plasmacytoid dendritic cells; pSTAT3: phosphorylated signal transducers and activators of transcription 3*

### MyD88 signaling trajectories differentiate patients with and without POCD

In contrast to the time-dependent differences observed for JAK/STAT signaling responses, other immune features, such as those contained in cluster 6, differed at all time points between the two patients’ groups, including prior to surgery (D0). Cluster 6 features were predominantly elements of the MyD88 signaling pathways in myeloid immune cell subsets, including cMCs, myeloid dendritic cells (mDCs) and plasmacytoid (p)DCs and in natural killer (NK) cells subsets (CD56^bright^CD16^neg^ NK cells). Notably, lower IκB signals in cMC, pDC, and mDC cells were observed at all time points in patients who developed POCD compared to controls (p = 0.032, 0.0041, and 0.0042 respectively, ANOVA, fixed effect for group, **Fig. 4B** and **Table S4**). Other time-independent differences in patients with POCD compared to controls included lower IκB signal in memory and naive B cells (p = 0.039 and 0.043 respectively, ANOVA, fixed effect for group) and lower B cell frequencies (p = 0.036 and < 0.001 respectively, ANOVA, fixed effect for group).

### A predictive model integrating preoperative single-cell and plasma proteomic immune responses accurately identifies patients at risk for POCD before surgery

The analysis of immune cell trajectories distinguishing patients with and without POCD highlighted postoperative immune cell features that may contribute to the pathogenesis of POCD. From a clinical perspective, the ability to differentiate patients’ immune states before surgery, in addition to the early post-operative period, is of paramount importance, as it allows for the implementation of effective, personalized preventive strategies. Remarkably, differences in cell-specific signaling pathways, including elements of the MyD88 signaling pathway in myeloid cell subsets, were observed prior to surgery (**Fig. 4**). Based on these findings, we broadened the immune profiling of patients’ samples collected before surgery with the goal of predicting later development of POCD. The mass cytometry immunoassay was expanded to include preoperative assessment of immune cell signaling activities in response to lipopolysaccharide (LPS) and CpG, agonists of TLR4 and TLR9, respectively, and activators of the MyD88 signaling pathway. The analysis of preoperative samples also included the concentration of 997 plasma proteins, quantified using the Somalogic Proteomics Assay(*40*) allowing integration of single-cell immune responses within the larger inflammatory proteomic network.

To identify preoperative single-cell or plasma proteomic features predictive of POCD, we employed Stabl(*41*), a sparse machine learning method that combines multivariable predictive modeling with a data-driven feature selection process. This method is optimized for analysis of multi-omic datasets, as each omic data layer is first examined individually prior to integration into a unique predictive model. In our study, the single-cell and plasma proteomic datasets were analyzed along with clinical and demographic data layers that may contribute to differences in patients’ risk for POCD, including hypertension, age, sex, surgery type, history of neurodegenerative disease, Charlson comorbidity index, level of education, type of anesthesia, fasting duration, preoperative cognitive, depressive, and anxiety status (**Fig. 5A**, **Table S5**). The model performance in predicting POCD was assessed using an external two-step cross-validation loop that included feature selection and model fitting. The analysis was performed using samples from a sub-cohort of 22 patients including 8 with POCD for which the complete single cell, proteomic, and clinical datasets were available. The analysis yielded a model accurately classifying patients with and without POCD (AUC = 0.80 [0.54, 0.9], p = 2×10^-2^ U-test, **Fig. 5B**). The multivariable predictive model comprised eleven features: ten mass cytometry features including eight intracellular signaling responses and two cell frequencies, as well as one proteomic feature **(Fig. 5C-F** and **Table S6)**, highlighting preoperative features predisposing patients to POCD.

**Figure 5:**
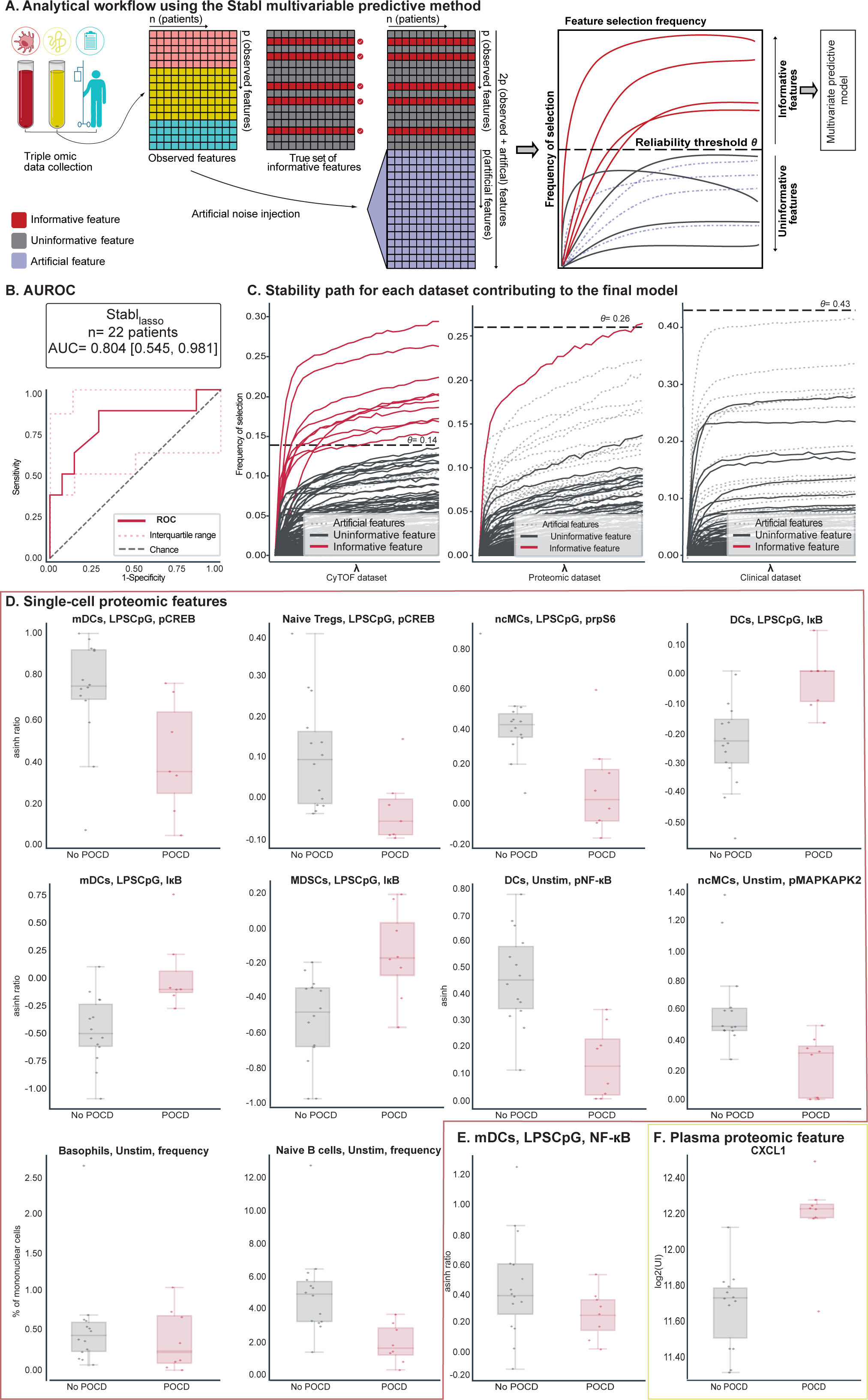
Multivariable modeling of preoperative single-cell and plasma proteomic and clinical datasets with data-driven selection of predictive biomarkers of POCD. **A**. Analytical workflow using the Stabl multivariable predictive modeling method (see methods). Model features are objectively selected using an artificial noise-injection technique allowing for data-driven estimation of a reliability threshold (theta) that controls the false discovery rate of feature selection. **B.** Predictive performance of the Stabl multivariable model for identifying patients who will develop POCD at POD7 (AUC = 0.80 [0.54, 0.9], p = 2×10^-2^ U-test, Monte Carlo cross-validation on a subset of 22 fully characterized subjects). **C.** Stability path showing data-driven feature selection for each dataset in the final model (*from left to right*: mass cytometry, plasma proteomics, clinical data). The horizontal dashed lines represent the reliability threshold, θ, specific to each dataset. The final model contains one plasma proteomic and ten single cell mass cytometry features. **D.** Barplots depicting the median and interquartile range of the single-cell mass cytometry features selected by the Stabl model and **E.** the median and interquartile range of NF-κB expression in mDCs, concomitant with the increase in IκB expression described in (A). **F.** Barplots depicting the median and interquartile range of the proteomic feature selected by the Stabl model. *Abbreviations: DCs: Dendritic cells; i*κB *inhibitor of factor kB; LPSCpG: blood sample stimulated with LPS & CpG; mDCs: myeloid dendritic cells; MDSCs: myeloid derived suppressor cells; ncMCs: non classical monocytes*; *NF-* κ*B: nuclear factor* κ*B POCD: post-operative cognitive decline; Tregs: Regulatory T cells; unstim: unstimulated blood sample*.

Among the most informative features of the predictive model were several elements of the MyD88 signaling pathway measured in myeloid immune cell subsets, including the IκB, pCREB, and prpS6 signals in mDCs, ncMCs, and myeloid derived suppressor cells (MDSCs). Notably, the IκB signal was dampened in DCs from patients who developed POCD (8.65 –log2 fold difference in median expression), particularly in mDCs (2.03 –log2 fold difference in median expression), and in MDSCs (1.37 –log2 fold difference in median expression). These results resonate with differences in the endogenous myeloid cell IκB signal observed after surgery and indicate that *ex vivo* TLR4/9 stimulation of samples before surgery recapitulates aspects of the endogenous innate immune response to surgery associated with POCD. Consistent with dampened MyD88 signaling responses in innate immune cells, the pCREB signal in mDCs was also decreased in patients with POCD (0.73 –log2fold difference respectively). Of note, pCREB is a key player in synaptic plasticity, and decreased levels of pCREB correlate with worse cognitive performance in rats(*42*) and with neurodegenerative diseases in humans(*42*, *43*). Additional model features included prpS6 in ncMCs, pCREB signaling in naïve regulatory T cells in response to TLR4/9 stimulation, basal naive B cell and basophil frequencies, basal pMAPKAPK2 in ncMCs, and pNF-κB in mDCs (**Table S7)**. In addition, the model included a single plasma proteomic feature, chemokine CXCL1 (0.43 –log2 fold difference in median expression), a potent chemoattractant known to be implicated in neuro-inflammatory processes such as Alzheimer’s disease(*44*). Interestingly, the predictive model did not select clinical features, consistent with the inability of previous clinical studies to accurately predict POCD from preoperative clinical data(*10*).

## Discussion

We conducted an extensive analysis of the immune response to major orthopedic surgery with the aim of identifying immunological factors associated with POCD in patients older than 60 years of age. Analysis of immune cell composition and intracellular signaling response dynamics after surgery revealed distinct immunological trajectories differentiating patients who developed POCD from controls. Furthermore, a sparse multivariable model integrating clinical, proteomic, and single-cell mass cytometry data collected before surgery accurately identified patients at risk for developing POCD (AUC of 0.80). The predictive model of POCD contained 11 single-cell and plasma proteomic features. Notably, the model lacked clinical parameters, consistent with the limited performance (AUC < 0.7) of previous predictive models of POCD based on pre-or post-operative clinical factors(*10*). The results underscore the importance of integrating biological variables that capture the pathophysiology of cognitive impairments into accurate prognostic assays(*11*) that can be leveraged for the personalized management of patients at risk for POCD.

Two key findings specified immunological dysfunctions in patients who developed POCD: (i) early exacerbation of pro-inflammatory JAK/STAT signaling responses in both innate and adaptive cells and (ii) dampened MyD88 signaling responses in myeloid cell subtypes that were detectable before surgery and persisted for 90 days post-surgery. These results point at distinct immunological trajectories integral to the pathophysiology of POCD and build upon prior knowledge of immune mechanisms predisposing patients to acute alterations in cognition, such as postoperative delirium(*33*, *45*, *46*), or to delayed recovery(*35*, *47*, *48*). For example, increased STAT3 signaling activity (i.e., intracellular pSTAT3 signal) in CD4^+^ T cells and cMCs at POD1 emerged as particularly informative single-cell features associated with POCD. Dysregulation of the JAK/STAT pathway has previously been implicated in neuroinflammatory disease pathogenesis, triggering and polarizing myeloid and T cells to pathogenic phenotypes, resulting in an excess of inflammatory cytokines and neurodegeneration(*49*). Notably, prolonged elevation of STAT3 has been recognized as a key contributor to neurodegenerative processes like Alzheimer’s disease(*50*). A persistent maladaptive chronic inflammatory state is also seen in frail adults, characterized by high levels of proinflammatory cytokines such as IL-6(*51*, *52*), the canonical upstream activator of STAT3 signaling, evoking “inflammaging”(*53*), one of the most conserved biological characteristics of frailty(*54*).

In our cohort, we also observed the dampening of innate immune cell MyD88 signaling responses to *ex vivo* TLR4/9 stimulation in patients who later developed POCD. This phenomenon affected key elements of the mitogen-activated protein kinase (MAPK), including MAPKAPK2, rpS6, and CREB, and the NF-κB (IκB) branches of the MyD88 pathways. Interestingly, the dampened MyD88 responses, particularly NF-κB signaling, is a hallmark of immunosenescence(*53*),(*55*), a dynamic and multifactorial process affecting both innate and adaptive immune cells, an important marker of inflammaging. Our findings align with previous studies investigating disparities in immune cell MyD88 signaling between frail patients and age-matched controls(*56–58*) and suggest a potential overlap in MyD88-dependent immunosenescence traits between patients with POCD and frail individuals(*56*). In addition, patients at risk for POCD exhibited elevated plasma CXCL1 concentrations. CXCL1 is a chemokine involved in microglia activation(*59*) and plays an important role in cell death and immunosenescense(*60*), with increased concentrations both causing and resulting from senescence(*60*). Together, the increased CXCL1 expression before surgery and dampened MyD88 innate immune cell responsiveness suggest that patients with POCD are in a state of chronic inflammation and immunosenescence, which may result in a maladaptive response to surgical stress contributing to the development of POCD.

This study has certain limitations. First, the cohort was assembled from an interventional trial investigating the effect of pre-operative ketamine administration on POCD after major orthopedic surgery. However, in our study, there was no difference in ketamine administration between the POCD and control groups and no effect of ketamine on POCD, preventing this variable from confounding our immunological analysis. Future studies in a larger population undergoing a broader variety of surgical interventions will be required to establish the generalizability of our results. Second, our assessment of immune signaling responses before surgery was limited to TLR signaling. In the future, it will be important to expand the panel of *ex vivo* stimulations. For example, including stimulation of JAK/STAT3 signaling responses (*e.g.*, using IL-6, IL-10, and IL-21 family members(*61*)) will be informative and may further improve the performance of our predictive model of POCD. Third, while mass cytometry allows for simultaneous detection of up to 50 parameters at a single-cell level, the technology necessitates the pre-selection of cell surface and intracellular features. Similarly, the proteomic platform included a selected panel enriched for inflammatory mediators. Future studies including untargeted transcriptomic, proteomic, or metabolomic approaches will help to ascertain whether a more comprehensive profiling of patients’ inflammatory, metabolic, and hormonal responses to traumatic injury can enhance the predictive power of these models.

In summary, the longitudinal mass cytometry assessment of systemic immune responses to surgery identified patient-specific immune trajectories that tracked the development of POCD. The findings pinpoint the roles of the JAK/STAT and MyD88 signaling pathways in the human pathophysiology of POCD. In addition, analysis of preoperative immune responses yielded a robust predictive model of POCD comprising a sparse number of immunological features detectable before surgery. The results hold promise for the development of clinically applicable predictive algorithms that can serve to facilitate an individual patient’s risk stratification and allocation of tailored care pathways to prevent POCD.

## Materials and Methods

### Experimental Design

This study was nested within the larger “Preoperative ketamine administration for prevention of postoperative cognitive decline after major orthopedic surgery in elderly patients” (POCK) study, a multicenter, randomized, placebo-controlled, quadruple-blind, superiority trial (ClinicalTrials.gov <u>NCT02892916</u>). After approval by the institutional review board (Comité de Protection des Personnes Ile de France III, 08/11/2016, France – #2016-000691-16) and obtention of written and signed consent, 33 patients aged 60 years and older, scheduled to undergo major orthopedic surgery under general anesthesia (e.g., spinal surgery including spinal decompression, hip or knee total joint arthroplasty, hip internal fixation, or shoulder arthroplasty) were enrolled at five university hospital centers in the Paris region between March 20, 2017, and May 28, 2019 (Hôpital Européen Georges Pompidou, Hôpital Lariboisière, Hôpital Pitié Salpêtrière, Hôpital Saint Antoine from Assistance Publique – Hôpitaux de Paris and Hôpital Foch). Exclusion criteria were an American Society of Anesthesiology score > 4, emergency surgery (i.e., emergency hip fracture), an expected length of stay in hospital < 48 hours, known allergy or contraindication to ketamine, severe auditory or vision disorders, drug misuse history or chronic anti-psychotic medications, severe alcoholic liver disease, or lack of French language fluency.

### Assessment of POCD

The primary clinical outcome of the study was the development of POCD on POD7. Cognition was assessed by trained members of the research team, using the MoCA test(*62*) and Trail Making Test (TMT) A and B, a validated tool to measure working memory (TMT A) and central executive functioning (TMT B)(*63*). An absolute difference in Z-score ≥ 1 in at least one cognitive test (MoCA or TMT A or TMT B or TMT B-A) between the tests carried out before surgery (D0) and on POD7, defined POCD(*64*), aligning the objective criteria of perioperative cognitive decline as described in DSM-5(*65*).

Other clinical data were assessed. Incidence of anxiety (defined as an anxiety score of 8 or above) and depression (defined as a depression score of 8 or above) were assessed using the Hospital and Anxiety Depression Scale(*66*) at POD7 and POD90; incidence of POCD, assessed by the same cognitive tests (MoCA or TMT A or TMT B or TMT B-A), three months after surgery at the surgical follow-up visit was also collected. (*62*, *63*) Pre– and postoperative pain was assessed by a patient-reported Visual Analog Scale at rest, from two hours after the end of surgery to POD7 on a daily basis and three months after surgery; cumulative opioid requirements were measured from the day of surgery (D0) to POD7.

## Single-cell and plasma proteomic analysis

### Blood collection

Whole blood, plasma samples, and clinical data were collected on D0 before induction of anesthesia and on three additional timepoints after surgery (POD1, 7, and 90). Whole blood samples were collected in sodium-heparinized tubes. They were then either left unstimulated to measure endogenous intracellular activities or exposed to a combination of the TLR4 agonist LPS and TLR9 agonist CpG. Only samples collected before surgery were stimulated. All samples were then fixed with Proteomic Stabilizer in Smart Tubes (Smart Tube Inc., San Carlos, CA) and immediately stored at ^−^80[°C. All samples were sent on dry ice to Stanford University (Stanford, CA).

### Barcoding, antibody staining, and mass cytometry processing

The 39-marker antibody panel consisted of 25 antibodies targeting cell surface markers and 14 intracellular antibodies specific to phosphorylated signaling epitopes. (**Table S7**). In brief, the samples were thawed, and red blood cells were lysed. Subsequently, the samples were barcoded and stained with both surface and intracellular antibodies following standardized protocols (*26*, *67*). To minimize variability between measurements, samples from each individual patient were barcoded, stained, and processed simultaneously on the CyTOF. To enhance the assay’s sensitivity for detecting differences between patients with POCD seven days after surgery and those without, time series samples from patients with POCD were randomly matched with samples from patients without POCD. These paired time series samples were then barcoded and processed using the same barcode plate. The barcoded samples were analyzed at a flow rate of 600 to 800 events per second. The output FCS files were normalized and de-barcoded using MatLab-based software. The FCS files were then uploaded to the Cell Engine (https://cellengine.com, Primity Bio, Fremont, CA) platform for gating.

### Cell frequency, endogenous intracellular signaling, and intracellular signaling responses

From each sample, 488 single-cell proteomic features, including the frequency of 31 major innate and adaptive immune cells defined using manual gates (**Fig. S2**) and their intracellular signaling activities (e.g., the phosphorylation state of 11 proteins including phospho-(p)STAT1, pSTAT3, pSTAT5, pSTAT6, pNF-κB, pMAPKAPK2, pP38, prpS6, pERK1/2, pCREB, and total IκBα) were analyzed. Immune cell frequency features were calculated for each immune cell subset from the unstimulated samples. For each cell type, intracellular activities were expressed as the median signal intensity (arcsinh transformed value). Signaling changes in response to receptor-specific ligands were reported as the arcsinh transformed ratio over the endogenous signaling, i.e., the difference in arcsinh transformed signal intensity between the stimulated and unstimulated condition. A penalization matrix, based on mechanistic immunological knowledge, was applied to signaling changes in response to receptor-specific ligand data. The proportion of mononuclear cells was expressed as a percentage of the gated live mononuclear cells (DNA positive and cPARP, CD235a, CD61, and CD66 negative). The proportion of neutrophils was expressed as a percentage of the gated live leukocytes (DNA positive and cPARP, CD235a, and CD61 negative).

### Plasma protein profiling

In parallel, the preoperative plasma concentrations of 997 proteins were quantified using a highly multiplex aptamer-based platform, the SOMAscan manual assay (version 1.4k) for human plasma. All analyses were performed in randomly allocated samples by SomaLogic, Inc. (Boulder, CO)(*68*). The assay quantifies relative concentrations of proteins over a wide dynamic range (>8 log) using chemically modified aptamers with slow off-rate kinetics (SOMAmer reagents). Each SOMAmer reagent is a unique, high-affinity, single-strand DNA endowed with functional groups mimicking amino acid side chains. Nucleotide signals are quantified using relative fluorescence on microarrays. The assay has a historic median intra– and inter-run coefficient of variation of around 5%, and median lower and upper limits of quantification of 3.0[pM and 1.5 nM(*40*). Relative levels of plasma proteins are reported in arbitrary units calculated from data normalized to internal controls and reported after log2 transformation.

## Statistical methods

### Preprocessing

Prior to statistical analysis, some initial preprocessing steps were applied to the acquired dataset. A knowledge-based penalization matrix was applied to intracellular signaling response features in the mass cytometry data based on mechanistic immunological knowledge, as previously described(*30*, *69*). Mechanistic priors used in the penalization matrix are independent of immunological knowledge related to POCD. Features with no variance were also removed from the dataset.

### Visualization, clustering, and longitudinal analysis of the single cell proteomic dataset

A correlation network using a t-distributed Stochastic Neighbor Embedding (t-SNE) model was performed for dimensional reduction and visualization of the single-cell proteomic dataset. We computed the correlation matrix using all the samples collected and all the available features within the unstimulated whole blood single cell panel. Edges are shown for correlations with a Pearson ρ> 0.9. The size of the nodes corresponds to –log(p-value) (Mann-Whitney rank-sum test) between patients with and without POCD.

### Clustering

Features were then clustered using the unsupervised framework k-means and the number of clusters was chosen using the silhouette score and optimizing it, looking at a range between four and 14 clusters, reaching a maximum at ten. We defined meta features representing the median of all features within a given cluster, to capture differences in the general trajectory of individual clusters between the two patients’ groups.

### Statistical tests

Original features and meta features were analyzed with a two-way ANOVA to estimate how the mean of each variable changes according to the levels of the two categorical variables: time-point and POCD status. All features at all timepoints were also compared using a Mann-Whitney U test. All analyses were performed using python3®. Libraries used included numpy, pandas, networks, matplotlib.pyplot, seaborn, scipy and sklearn.

### Power analysis

A power analysis was performed to determine the required sample size based on the following parameters: an estimated AUROC (θ) of 0.90, indicating a high level of predictivity; a proportion (P) of 0.40, representing the expected prevalence of POCD within our cohort; a confidence interval width (W) of 0.25; and a confidence level (CL) of 90%. The resultant sample size required for our study was determined to be 23 patients (*70*).

### Stabl, predictive modeling

The Stabl algorithm was employed to identify features predictive of POCD. Stabl is a supervised machine learning framework that allows for sparse, reliable, and predictive feature selection. Prior to model fitting, the preprocessed features underwent z-scoring.

As previously described,(*71*) Stabl fits sparsity–promoting regularization methods (e.g., Lasso or the elastic net) on data subsamples using a procedure similar to Stability Selection. Subsampling mimics the availability of multiple random cohorts and estimates each feature’s selection frequency across all iterations.

To determine the optimal frequency threshold, Stabl introduces artificial features unrelated to the outcome (noise injection) via random permutations. The assumption is that these artificial features behave similarly to uninformative features in the original dataset. The artificial features are used to construct a surrogate of the false discovery proportion (FDP_+_). We define the “reliability threshold”, θ, as the frequency threshold that minimizes FDP_+_ across all possible thresholds. This objective, data-driven approach to establishing θ adapts to individual omic datasets. The set of features with a selection frequency larger than θ (i.e., reliable features) is included in a final predictive model. The complete package for Stabl is available online at https://github.com/gregbellan/Stabl. Features were selected from three datasets (single cell and plasma proteomics and clinical data) separately and merged into a unified model.

### Cross validation

To assess our model’s predictive performance, we utilized a Monte Carlo cross-validation approach tailored for the integration of multi-omic data modalities, restricting our analysis to a subset of 22 fully characterized subjects with complete single-cell, proteomic, and clinical data available. At each fold, the dataset is split randomly into training and testing sets, and the model is then trained and evaluated using the training and testing sets respectively. The cross-validation procedure was performed using the Repeated Stratified K-Fold class of ‘scikit-learn’ (v1.1.2) which repeats multiple K-fold cross-validation schemes. We then take the median of the predictions to obtain the final predictions. This technique ensures that all samples are evaluated the same number of times. We used stratified 5-fold cross-validation (20% of the data is tested at each fold) to ensure that the class repartition was preserved among all the folds.

The code and the data are documented and available at https://github.com/gregbellan/POCD.

## Acknowledgments

None

## Data and materials availability

All data are available in the main text, the supplementary materials or at https://github.com/gregbellan/POCD.

## Author contributions

Conceptualization: FV, BG

Methodology: FV, BG, JH

Investigation: FV, BG, JH, EG, IAS, DF, AB, KA, JE, BC, MS

Visualization: FV, AC

Funding acquisition: FV, BG

Project administration: BG

Supervision: BG, BH, DD, RG, TS, SM, NA, MA

Writing – original draft: FV, JH, AC, GB

Writing – review & editing: DG, BG

