## Supplementals for "An immune signature of postoperative cognitive decline in elderly patients"

Franck Verdonk *et al*

**This PDF file includes:**

Figs S1 to S4

Tables S1 to S8

**Fig. S1.**

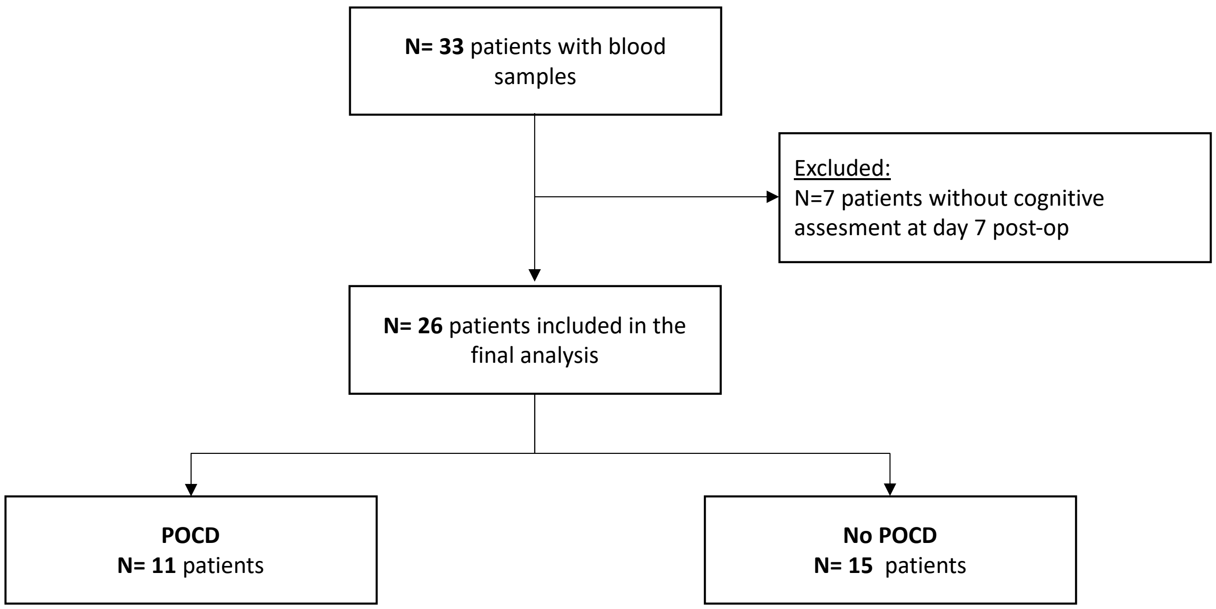


**Figure S1:** Study flow chart according to CONSORT guidelines. *Abbreviations: POCD: Post-operative cognitive dysfunction.*

**Fig. S2.**

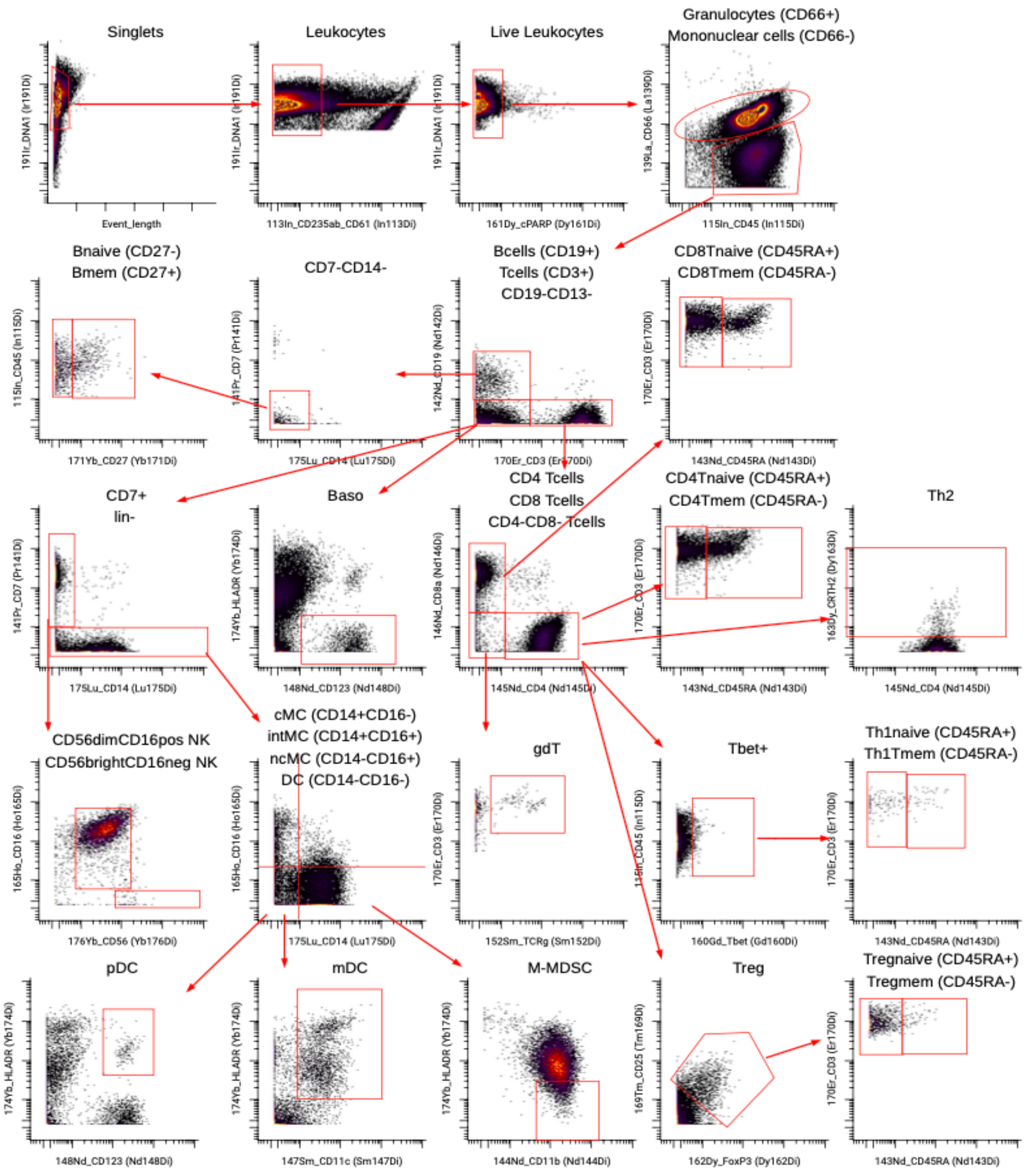


**Figure S2:** Immune cell gating strategy. Gating strategy for the identification of immune cell subsets included in the analysis.

### Fig. S3.

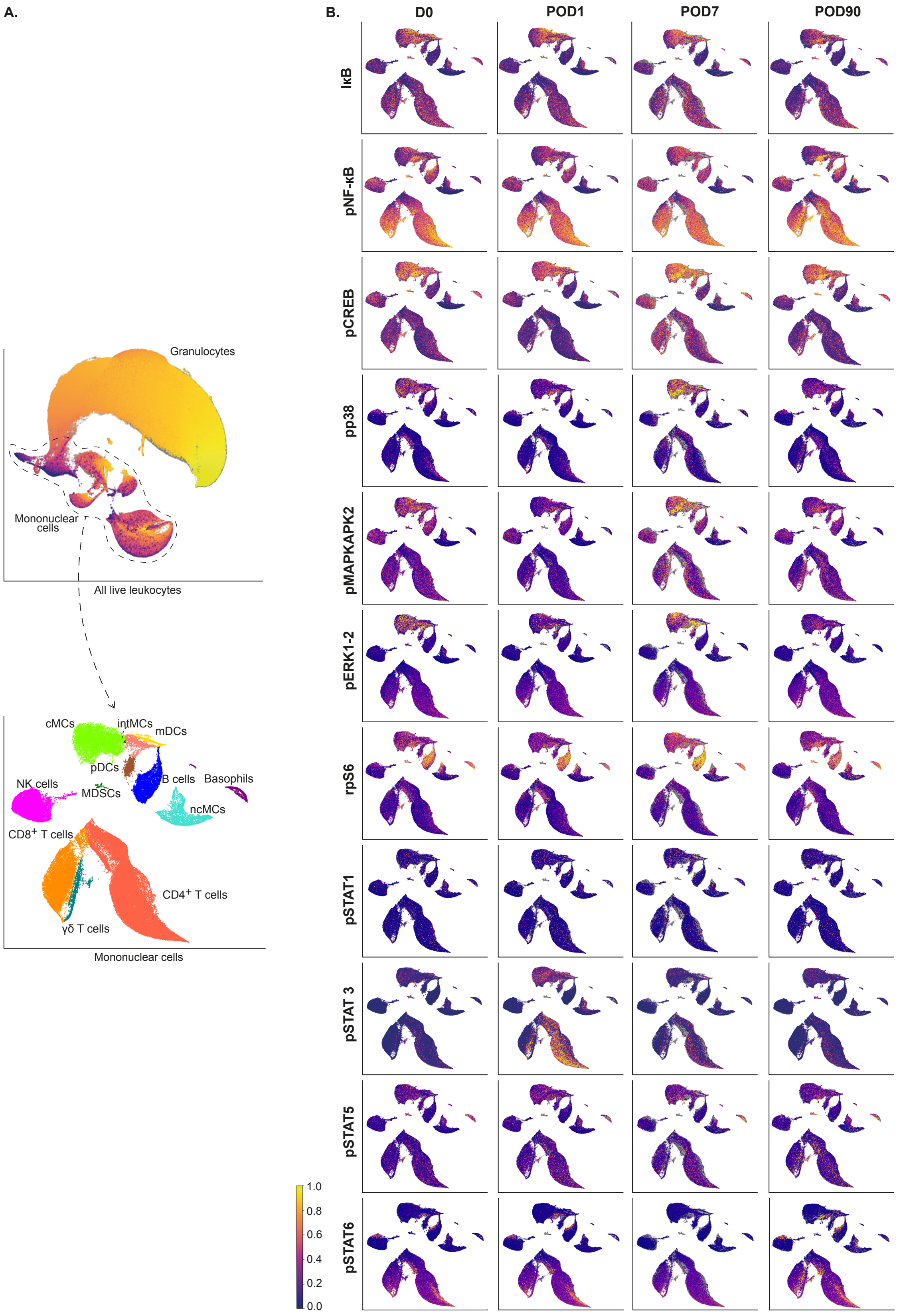
A.
Granulocytes
Mononuclear cells
All live leukocytes
cMCs
intMCs
mDCs
pDCs
B cells
Basophils
NK cells
MDSCs
ncMCs
CD8+ T cells
CD4+ T cells
γδ T cells
Mononuclear cells
B.
D0
POD1
POD7
POD90
IκB
pNF-κB
pCREB
pp38
pMAPKAPK2
pERK1-2
rpS6
pSTAT1
pSTAT 3
pSTAT5
pSTAT6
1.0
0.8
0.6
0.4
0.2
0.0

**Figure S3 : A.** UMAP representation of the single-cell mass cytometry dataset. Top, all live leukocytes, including neutrophils and mononuclear cells; Bottom: UMAP representation of mononuclear cells only. UMAPS are clustered by cell types and annotated. **B.** UMAPs representing all mononuclear cells colored according intra-cellular signaling at D0 (before surgery), D1, D7 and D90 postoperative. *Abbreviations: cMCs: classical monocytes; CREB: cAMP-response element binding protein; D: Day; γδ T cells: gamma delta T cells; iκB: inhibitor factor κB; intMCs: intermediate monocytes; mDCs: myeloid dendritic cells; MDSCs: Myeloid Derived Suppressor Cells; NF-κB: Nuclear factor – κB; ncMCs: non classical monocytes; NK cells: natural killer cells; pDCs: plasmacytoid dendritic cells; p: phosphorylation; STAT: Janus Kinase/signal transducers and activators of transcriptions; rpS6 ribosomal protein S6.*

**Fig. S4.**

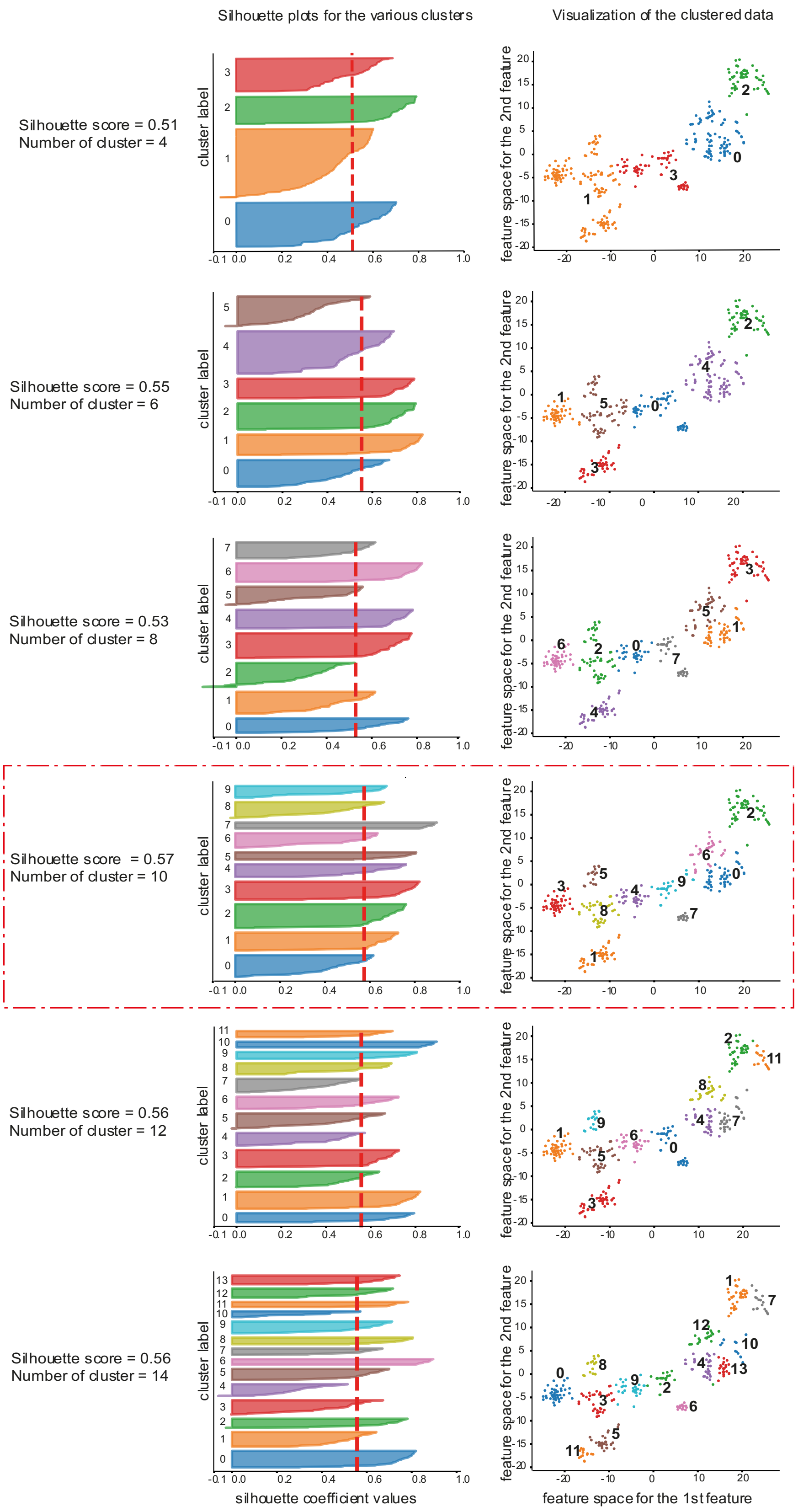
Silhouette plots for the various clusters
Visualization of the clustered data
Silhouette score = 0.51
Number of cluster = 4
Silhouette score = 0.55
Number of cluster = 6
Silhouette score = 0.53
Number of cluster = 8
Silhouette score = 0.57
Number of cluster = 10
Silhouette score = 0.56
Number of cluster = 12
Silhouette score = 0.56
Number of cluster = 14
cluster label
feature space for the 2nd feature
silhouette coefficient values
feature space for the 1st feature

**Figure S4:** Silhouette scores. Adequate number of cluster was chosen according to silhouette score. On the left are presented the silhouette scores for 4, 6, 8, 10, 12 and 14 clusters. The red dashed-line represents the average silhouette score. On the right, visualization of the clustered data on a t-SNE two-dimensional representation.Ten clusters is the most optimized number of clusters.

**Table S1.**

| **Characteristics** | **Whole population (n=26)** | **No POCD (n=15)** | **POCD (n=11)** | **p-value** |
|---|---|---|---|---|
| **Opioid use at anesthesia induction,** yes | 26 (100%) | 15 (100%) | 11 (100%) | 0.348 |
| **Per operative opioid use,** yes | 25 (96.15%) | 15 (100%) | 10 (90.91%) | 0.595 |
| **Dexamethasone use**, yes | 19 (73.08%) | 11 (73.33%) | 8 (72.73%) | 0.295 |
| **Perioperative use of ketamine,** yes | 11 (42.31%) | 7 (46.67%) | 4 (36.36%) | 0.295 |
| **Surgery duration,** in minutes | 157.5 [120 ;198] | 178 [130 ;215] | 120 [110 ;160] | **0.009** |

**Table S1**: Peri-operative management of the study population. Values are expressed as number (%). Both group did not differ in any drug use or surgery type. *Abbreviations: POCD: Post-operative Cognitive Disorder*

**Table S2.**

| **Characteristics** | **Whole population (n=26)** | **No POCD (n=15)** | **POCD (n=11)** | **p-value** |
|---|---|---|---|---|
| **Opioid use in the PACU**, yes | 26 (100%) | 15 (100%) | 11 (100%) | 0.348 |
| **Cumulative morphine dose from PACU to POD6,** median[IQR]* | 17 [10;42.5] | 14.5 [8.5;36.25] | 24 [15;45] | 0.407 |
| **Pain (VAS) in the PACU,** median [IQR]** | 4 [1;6] | 5 [2.25;5.75] | 4 [0;6.5] | 0.613 |
| **Pain (VAS) at POD1,** median [IQR]** | 3 [2;5] | 3 [2.25;4] | 3 [2.5;5] | 0.362 |
| **Pain (VAS) at POD7,** median [IQR] | 1[0.5 ;5] | 2 [0.5;4.5] | 0 [0;5] | 0.685 |
| **Pain (VAS) at POD90,** median [IQR]*** | 2 [0;4] | 2 [0;4] | 2 [0;5.75] | 1 |
| **Anxiety and depression status** | | | | |
| Anxiety (HADS Anxiety score of 8 or above) at POD7† | 7 (30.43%) | 4 (28.57%) | 3 (33.33%) | 1 |
| Depression (HADS Depression score of 8 or above) at POD7† | 2 (8.7%) | 0 (0%) | 2 (22.22%) | 0.142 |
| Anxiety (HADS Anxiety score ≥ 8) at D90*** | 9 (40.91%) | 4 (33.33%) | 5 (50%) | 0.666 |
| Depression (HADS Depression score ≥ 8) at POD90*** | 4 (18.18%) | 2 (16.67%) | 2 (20%) | 1 |

**Table S2 :** Post-operative clinical assessment of the study population. Both groups did not differ in any category. Values are expressed as number (%) or median [IQR].

*Abbreviations: HADS: Hospital Anxiety and Depression Scale; IQR: Interquartile Range, MOCA; Montreal Cognitive Assessment Score; POCD: Post-operative Cognitive Disorder, POD: post-operative day*

* Denominator is 19 for *all groups*, 10 for *no POCD* and 9 for *POCD* groups

** Denominator is 25for *all groups*, 14 for *no POCD* and 11 for *POCD* groups

*** Denominator is 22 for *all groups*, 12 for *no POCD* and 10 for *POCD* groups

† Denominator is 23 for *all groups*, 14for *no POCD* and 9 for *POCD* groups

**Table S3 .**

| | Feature signaling type | Cluster 0 N= 52 | Cluster 1 N= 43 | Cluster 2 N=56 | Cluster 3 N=43 | Cluster 4 N=31 | Cluster 5 N=18 | Cluster 6 N=35 | Cluster 7 N= 15 | Cluster 8 N=38 | Cluster 9 N=28 |
|---|---|---|---|---|---|---|---|---|---|---|---|
| **MyD88** | **IκB** | | | | | 1 (3.2%) | | 24 (68.6%) | | 3 (7.9%) | 2 (7.1%) |
| | **NF-κB** | | | 25 (44.6%) | | | | 1 (2.9%) | | 1 (2.6%) | 3 (10.7%) |
| | **pERK12** | 3 (5.8%) | | | 1 (2.3%) | 3 (9.7%) | | | | 8 (21.1%) | 15 (53.6%) |
| | **pP38** | 4 (7.7%) | 26 (60.5%) | | | | | | | | |
| | **pCREB** | 19 (36.5%) | | 5 (8.9%) | | | | 4 (11.4%) | | | 2 (7.1%) |
| | **pMAP-KAPK2** | 26 (50.0%) | 2 (4.6%) | | | 2 (6.5%) | | | | | |
| **JAK/STAT** | **pSTAT1** | | | | 30 (69.8%) | | | | | | |
| | **pSTAT3** | | | | 12 (27.9%) | | 18 (100%) | | | | |
| | **pSTAT5** | | | | | 5 (16.1%) | | | | 24 (63.2%) | 1 (3.6%) |
| | **pSTAT6** | | 15 (34.9%) | | | 18 (58.1%) | | | 15 (100%) | | |
| **prpS6** | | | | 5 (8.9%) | | | | 4 (11.4%) | | | 3 (10.7%) |
| **Frequency** | | | | 21 (37.5%) | | 2 (6.5%) | | 2 (6.5%) | | 2(5.3%) | 2 (7.1%) |

**Table S3:** Intracellular signaling pathway in each cluster. Number of features belonging to each signalling pathway is expressed as number (%).

Table S4.

| Feature | POCD | Time | POCD : Time | Cluster |
|---|---|---|---|---|
| **B cells, Frequency** | 0.00014 | 0.96789 | 0.97223 | 2 |
| **B naive, Frequency** | 0.00015 | 0.78260 | 0.81579 | 2 |
| **NK, Frequency** | 0.01795 | 0.13025 | 0.83029 | 2 |
| **$CD56^{dim}CD16^{pos}$NK, Frequency** | 0.03445 | 0.09510 | 0.95041 | 2 |
| **Th1, pERK12** | 0.04917 | 0.17630 | 0.19348 | 2 |
| **pDC, pSTAT1** | 0.01777 | 0.87558 | 0.85247 | 3 |
| **Memory B cells, IkB** | 0.03912 | 0.28247 | 0.89280 | 4 |
| **gdT cells, pSTAT3** | 0.03179 | 0.13069 | 0.03635 | 3 |
| **Treg naive, pSTAT3** | 0.00021 | 7.38501E-15 | 0.07031 | 5 |
| **CD4T cells naive, pSTAT3** | 0.00465 | 5.37213E-16 | 0.23877 | 5 |
| **CD4T cells, pSTAT3** | 0.00792 | 3.79358E-11 | 0.30885 | 5 |
| **memory CD4T cells, pSTAT3** | 0.00800 | 1.99246E-10 | 0.33259 | 5 |
| **cMC, pSTAT3** | 0.01080 | 0.00729 | 0.46300 | 5 |
| **Treg, pSTAT3** | 0.01252 | 1.77776E-10 | 0.18490 | 5 |
| **Memory Treg, pSTAT3** | 0.01335 | 1.57405E-10 | 0.21072 | 5 |
| **Granulocytes, pSTAT3** | 0.01428 | 0.00008 | 0.15004 | 5 |
| **T cells, pSTAT3** | 0.03490 | 1.44165E-07 | 0.50586 | 5 |
| **ncMC, pSTAT3** | 0.03819 | 0.00010 | 0.02033 | 5 |
| **intMC, pSTAT3** | 0.04659 | 0.00034 | 0.04781 | 5 |
| **mDC, IκB** | 0.00412 | 0.09509 | 0.14135 | 6 |
| **pDC, IκB** | 0.00428 | 0.21415 | 0.78866 | 6 |
| **cMC, IκB** | 0.03280 | 0.14012 | 0.85675 | 6 |
| **Memory B cells, Frequency** | 0.03640 | 0.78591 | 0.92584 | 6 |
| **DC, IκB** | 0.03653 | 0.25451 | 0.22305 | 6 |
| **$CD56^{bright}CD16^{neg}$NK, IkB** | 0.04424 | 0.01023 | 0.62459 | 6 |
| **Naive Treg, pSTAT5** | 0.01738 | 0.06987 | 0.03408 | 8 |
| **$CD56^{bright}CD16^{neg}$NK, pERK1-2** | 0.01774 | 0.73636 | 0.59402 | 8 |
| **Naive B cells, IκB** | 0.04289 | 0.25730 | 0.68070 | 8 |
| **Naive Treg, pERK1-2** | 0.00344 | 0.64727 | 0.15509 | 9 |

**Table S4 :** Key intracellular features of preoperative unstimulated blood samples features. Two-way analyses of variance (ANOVAs) cognitive status x timepoint were performed and 29 top intra-cellular features were identified. The "POCD" and "Time" columns indicate if the cognitive status (no POCD or POCD) or the timepoint (D0, POD1, 7 and 90) could by itself affect the results. The POCD:Time column indicates if an interaction between the condition (no POCD or POCD) and the timepoint (D0, POD1, 7 and 90) could affect the results. The features are grouped into clusters as shown in the "Cluster" column.

**Table S5.**

| **Preoperative clinical features** | **Preoperative clinical features** |
|---|---|
| **Randomization arm** | Preoperative Morphine consomption |
| **Sexe** | Preoperative delirium (CAM) |
| **Age** | Preoperative MOCA score |
| **Weight** | Preoperative TMT-A score |
| **Height** | Preoperative TMT-B score |
| **BMI** | Preoperative HADS anxiety score |
| **Level of education** | Preoperative HADS depression score |
| **Comorbidities*** | Preoperative Apathy score |
| **Medication load, number** | Preoperative Starkstein score |
| **Chronic medication, type **** | Preoperative Charlson score |
| **Electrolyte disorder** | Surgery type |
| **Anemia** | Date of surgery |
| **Active smoking** | Anesthesia type (general or medullar) |
| **Preoperative pain (VAS)** | Regional anesthesia |
| **Neuropathic pain** | Fasting duration |

**Table S5 :** Clinical features used to build the Stabl model

*Comorbidities include (alphabetical order): Asthma, Cancer, Chronic pulmonary disease, Dementia, Diabetes, Epileptia, Hemiplegia, Heart failure, HIV stage AIDS, Hypertension, Ischemic cardiopathy, Leukemia, Liver disease, Lymphoma, Metastatic Cancer, Obstructive sleep apnea, Parkinson disease, Renal failure, Stroke, Vascular disease,

** Prescription drugs recorded and included in the model are : ACE inhibitors, antiplatelets agents, anti-viral medication, betablockers, low molecular weight heparin, lansoprazole, methotrexate, non-steroidal anti-inflammatory drugs, oral antidiabetic, oral direct anticoagulants.

Abbreviations : BMI : Body mass index; VAS: Visual analogical scale

**Table S6.**

| Feature | Associated weight |
| --- | --- |
| **ncMCs, Unstim, pMAPKAPK2** | -1.6624374101052044 |
| **DCs, Unstim, pNF-κB** | -2.374897952417473 |
| **Basophils, Unstim, Frequency** | -0.19775601782629854 |
| **Naive B cells, Unstim, Frequency** | -5.381525546166335 |
| **mDCs, LPSCpG, pCREB** | -0.7836505748272852 |
| **Naive Tregs, LPSCpG, pCREB** | -2.6190926984260154 |
| **MDSCs, LPSCpG, IκB** | 1.9150433973839314 |
| **DCs, LPSCpG, IκB** | 1.695597161776877 |
| **mDCs, LPSCpG, IκB** | 3.77712497219793 |
| **ncMCs, LPSCpG, prpS6** | -1.1913902099195244 |
| **CXCL1** | 4.7734176093927045 |

**Table S6:** Predictive Stabl model features and their associated weight. Features were selected from both single cell and plasma proteomic dataset using the Stabl algorithm. Their relative weight in the prediction model is displayed in the second column. Features are labelled by cell type, stimulation, signaling pathway (or frequency), for single cell proteomic features and by name for plasma proteomic features. *Abbreviations: DC: Dendritic cells; inhibitor of factor kB; LPSCpG: blood sample stimulated with LPS & CpG; mDC: myeloid dendritic cells; MDSC: myeloid derived suppressor cells; ncMC: non classical monocytes; POCD: post-operative cognitive decline; Treg: Regulatory T cells; unstim: unstimulated blood sample*

**Table S7.**

| Feature | Fold difference in median expression |
|---|---|
| **ncMCs, Unstim, pMAPKAPK2** | 0.98 -log2 |
| **DCs, Unstim, pNF-κB** | 0.46 -log2 |
| **Basophils, Unstim, Frequency** | 1.02 -log2 |
| **Naive B cells, Unstim, Frequency** | 0.60 -log2 |
| **mDCs, LPSCpG, pCREB** | 0.73 -log2 |
| **Naive Tregs, LPSCpG, pCREB** | 1.79 -log2 |
| **MDSCs, LPSCpG, IκB** | 1.37 -log2 |
| **DCs, LPSCpG, IκB** | 8.65 -log2 |
| **mDCs, LPSCpG, IκB** | 2.03 -log2 |
| **ncMCs, LPSCpG, prpS6** | 0.31 -log2 |
| **CXCL1** | 0.43 -log2 |

**Table S7: fold difference in median expression of the model features between patients with and without POCD.** Features are labelled by cell type, stimulation, signaling pathway (or frequency), for single cell proteomic features and by name for plasma proteomic features. *Abbreviations: DC: Dendritic cells; inhibitor of factor kB; LPSCpG: blood sample stimulated with LPS & CpG; mDC: myeloid dendritic cells; MDSC: myeloid derived suppressor cells; ncMC: non classical monocytes; POCD: post-operative cognitive decline; Treg: Regulatory T cells; unstim: unstimulated blood sample*

**Table S8.**

| Antibody | Manufacturer | Metal | Isotope | Clone | Concentration | Comment |
|---|---|---|---|---|---|---|
| **Barcode1** | Trace Sciences | Pd | 102 | | 15µM | Barcode |
| **Barcode2** | Trace Sciences | Pd | 104 | | 15µM | Barcode |
| **Barcode3** | Trace Sciences | Pd | 105 | | 15µM | Barcode |
| **Barcode4** | Trace Sciences | Pd | 106 | | 15µM | Barcode |
| **Barcode5** | Trace Sciences | Pd | 108 | | 15µM | Barcode |
| **Barcode6** | Trace Sciences | Pd | 110 | | 15µM | Barcode |
| **CD235ab** | Biolegend | In | 113 | HIR2 | 1µg/mL | Phenotype |
| **CD61** | BD | In | 113 | VI-PL2 | 0.5µg/mL | Phenotype |
| **CD45** | Biolegend | In | 115 | HI30 | 1µg/mL | Phenotype |
| **CD66** | BD | La | 139 | CD66a-B1.1 | 0.5µg/mL | Phenotype |
| **CD7** | BD | Pr | 141 | M-T701 | 0.5µg/mL | Phenotype |
| **CD19** | Biolegend | Nd | 142 | HIB19 | 0.5µg/mL | Phenotype |
| **CD45RA** | Biolegend | Nd | 143 | HI100 | 0.5µg/mL | Phenotype |
| **CD11b** | Biolegend | Nd | 144 | ICRF44 | 2µg/mL | Phenotype |
| **CD4** | Biolegend | Nd | 145 | RPA-T4 | 2µg/mL | Phenotype |
| **CD8a** | Biolegend | Nd | 146 | RPA-T8 | 1µg/mL | Phenotype |
| **CD11c** | Biolegend | Sm | 147 | Bu15 | 1µg/mL | Phenotype |
| **CD123** | Biolegend | Nd | 148 | 6H6 | 1µg/mL | Phenotype |
| **pCREB** | Cell Signaling Technology | Sm | 149 | 87G3 | 2µg/mL | Function |
| **pSTAT5** | Cell Signaling Technology | Nd | 150 | C11C5 | 4µg/mL | Function |
| **pP38** | BD | Eu | 151 | 36/p38 | 2µg/mL | Function |
| **TCRgd** | BD | Sm | 152 | B1 | 4µg/mL | Phenotype |
| **pSTAT1** | BD | Eu | 153 | 14/P-STAT1 | 1µg/mL | Function |
| **pSTAT3** | Cell Signaling Technology | Sm | 154 | M9C6 | 2µg/mL | Function |
| **pS6** | Cell Signaling Technology | Gd | 155 | D57.2.2E | 2µg/mL | Function |
| **FceRI** | Biolegend | Gd | 156 | AER-37 (CRA-1) | 0.5 µg/mL | Phenotype |
| **CD33** | Biolegend | Gd | 158 | WM53 | 2µg/mL | Phenotype |
| **pMAPKAPK2** | Cell Signaling Technology | Tb | 159 | 27B7 | 1µg/mL | Function |
| **Tbet** | Thermo Fisher | Gd | 160 | 4B10 | 8µg/mL | Phenotype |
| **cPARP** | BD | Dy | 161 | F21-852 | 1µg/mL | Phenotype |
| **FoxP3** | Thermo Fisher | Dy | 162 | PCH101 | 8µg/mL | Phenotype |
| **IκB** | Cell Signaling Technology | Dy | 164 | L35A5 | 8µg/mL | Function |
| **CD16** | Biolegend | Ho | 165 | 3G8 | 1µg/mL | Phenotype |
| **pNF-κB** | BD | Er | 166 | K10-895.12.50 | 2µg/mL | Function |
| **pERK1-2** | Cell Signaling Technology | Er | 167 | D13.14.4E | 4µg/mL | Function |
| **pSTAT6** | Biolegend | Er | 168 | A15137E | 1µg/mL | Function |
| **CD25** | Biolegend | Tm | 169 | M-A251 | 2µg/mL | Phenotype |
| **CD3** | Biolegend | Er | 170 | UCHT1 | 1µg/mL | Phenotype |
| **CD27** | BD | Yb | 171 | M-T271 | 2µg/mL | Phenotype |
| **CD15** | Biolegend | Yb | 172 | W6D3 | 8µg/mL | Phenotype |
| **CCR2** | Biolegend | Yb | 173 | K036C2 | 2µg/mL | Phenotype |
| **HLA-DR** | Fluidigm | Yb | 174 | L243 | 2µg/mL | Phenotype |
| **CD14** | Fluidigm | Lu | 175 | M5E2 | 2µg/mL | Phenotype |
| **CD56** | BD | Lu | 176 | NCAM16.2 | 1µg/mL | Phenotype |
| **DNA1** | Fluidigm | Ir | 191 | | 50µM | DNA |
| **DNA2** | Fluidigm | Ir | 193 | | 50µM | DNA |

**Table S8:** Mass Cytometry antibody panel
